## supplementary material for "Disentangling the determinants of transposable elements dynamics in vertebrate genomes using empirical evidences and simulations"

Supplementary Methods

For the convenience of the reader, we reproduce below parts of Methods sections found in previous studies [1,2] describing 1) how we collected DNA and obtained sequencing data from anoles samples 2) how we estimated recombination rate along the genome from SNPs, 3) how regions under positive selection were identified based on SNP data.

**DNA Extraction and Whole Genome Sequencing**

Whole genome sequencing libraries were generated from *Anolis carolinensis* liver tissue samples collected between 2009 and 2011 [3], and *A. porcatus* and *A.* *allisoni* tissue samples generously provided by Breda Zimkus from the Museum of Comparative Zoology at Harvard University. For each of the 29 samples, DNA was isolated from ethanol preserved tissue using Ampure beads per the manufacturers protocol. Illumina TRU-Seq paired end libraries were generated using 200 ng of DNA per sample and sequenced at the NYUAD Center for Genomics And Systems Biology Sequencing Core (http://nyuad.nyu.edu/en/research/infrastructure-and-support/core-technology-platforms.html) with an Illumina HiSeq 2500. Read quality was assessed with FastQCv0.11.5 (http://www.bioinformatics.babraham.ac.uk/projects/ fastqc) and Trimmomatic [4] was subsequently used to remove low quality bases, sequencing adapter contamination and systematic base calling errors. Specifically, the parameters “trimmomatic_adapter.fa:2:30:10 TRAILING:3 LEADING:3 SLIDINGWINDOW:4:15 MINLEN:36” were used. Samples had an average of 1,519,339,234 read pairs, and after quality trimming 93.3% were retained as paired reads and 6.3% were retained as single reads.

**Sequence Alignment and SNP Calling**

Quality trimmed reads were aligned to the May 2010 assembly of the *A. carolinensis* reference genome (Broad AnoCar2.0/anoCar2; GCA_000090745.1; [5]) and processed for SNP detection with the assistance of the NYUAD Bioinformatics Core, using NYUAD variant calling pipeline. Briefly, the quality-trimmed FastQ reads of each sample were aligned to the AnoCar2.0 genome using the BWA-mem short read alignment approach [6]and resulting SAM files were converted into BAM format, sorted and indexed using SAMtools [7]. Picard was then used to identify insertions, deletions and duplications in the sorted BAM files (http://broadinstitute.github.io/ picard/) and evaluated using SAMtools (stats and depth). Alignments contained an average of 204,459,544 reads that passed QC, 97.75% mapping and 91.93% properly paired. Each individual re-sequenced genome was then processed with GATK for indel realignment, SNP and indel discovery and genotyping, following GATK Best Practices [8,9]. GATK joint genotyping was conducted with HaplotypeCaller for increased sensitivity and confidence, and results were selectively compared to results generated from SAMtools mpileup [7]. Filtering was performed in VCFtools [10], with the following criteria: a 6X minimum depth of coverage per individual, a 15X maximum average depth of coverage, no more than 40% missing data across all 29 samples, a minimum quality score of 20 per site, and a minimum genotype quality score of 20.

**Estimating recombination rates**

We used the LDHat software [11] to estimate effective recombination rates (*ρ*=4*N_e_r* with *r* the recombination rate per generation and *N* the effective population size) along the green anole genome. This method has been successfully used to obtain recombination maps for datasets similar to ours in terms of sequencing depth and sample sizes (e.g. [12]). Unphased genotypes were converted into LDHat format using VCFtools (option –ldhat). Since LDHat assumes that samples are drawn from a panmictic population, we focused on the Eastern Florida clade for which sampling effort was the highest (n=8 diploid individuals). We used precomputed likelihood lookup tables with an effective population mutation rate (*θ*) of 0.001, which was the closest from the *θ* value estimated from our dataset (*θ* ~ 0.004) and used the lkgen module to generate a table fitting the number of observed samples (16 chromosomes). Recombination rates were estimated over 500kb windows with 100kb overlaps using the Bayesian reversible MCMC scheme implemented in the interval module. The chain was run for 1,000,000 iterations and sampled every 5000 iterations with a large block penalty of 20 to avoid overfitting and minimize random noise. The first 100,000 generations were discarded as burn-in. Convergence under these parameters was confirmed by visually inspecting MCMC traces for a subset of windows. We averaged *ρ* estimates over non-overlapping 1Mb windows along the genome.

**Genome scan using diploS/HIC**

We used diploS/HIC, a recently developed machine-learning algorithm, to classify genomic windows as selected or not in northern clades. Simulated datasets are split into subwindows that are described by a set of 12 summary statistics [13] recapitulating the allele frequency spectrum or linkage disequilibrium. In the case of selection, simulated windows where the selected site lies in the central subwindows are considered hard or soft sweeps examples, while other windows are considered linked-hard or linked-soft examples. While hard sweeps correspond to events where a new mutation is immediately advantageous and rises in frequency, soft sweeps take place when an advantageous mutation has been present in a population for long enough and recombined before selection took place, or when several advantageous mutations appear at the same locus. In that case, several different haplotypes may increase in frequency. Such events (‘soft sweeps’), as well as partial hard sweeps, are less efficiently detected by GWSS based on cross-population comparison [14], as they do not erase polymorphism to the same extent as completed hard sweeps. This set of simulated datasets is then used to train a supervised machine-learning algorithm that uses the spatial organization of summary statistics to differentiate between each category. Predictions on the actual genomic dataset are then performed over subwindows along the genome using the trained model. The procedure is similar in spirit to Approximate Bayesian Computation (ABC, [15,16]), where a set of simulated summary statistics is compared to an observed dataset to infer relevant population genetics parameters. However, machine learning is less sensitive to the choice of summary statistics, and requires less simulations [17].

We followed a procedure similar in spirit to a previous study [18]. We trained the algorithm using a set of 3,000 coalescent simulations using the discoal simulator [19], modelling changes in population sizes inferred from a previous SMC++ [20] analysis (see [2] and Sup Figure 1, this study). This coalescent analysis based on whole-genome sequences showed that Northern populations (CA and GA) displayed a clear signature of expansion starting between 200,000 and 100,000 years ago, following a bottleneck that started between 500,000 and 1,000,000 years in the past. We simulated 330kb windows divided into 11 subwindows. Hard and soft sweep examples consisted in windows with a sweep occurring in the central 30kb subwindow. We sampled selection coefficients (*s*) from a uniform distribution so that the product with the effective population size (*N*), 2*Ns*, covered the range (30, 3000) for soft sweeps and 2*Ns* ~ (30,300) for hard sweeps. This choice of different ranges was due to the stronger effect of hard sweeps that erased all diversity over the simulated window above 2*Ns*=300. For soft sweeps, we used an uniform distribution on the initial frequency of the adaptive variant covering the range (0,0.2). This set of parameters was chosen after multiple pilot tests to be as broad as possible while retaining power to detect selective events. We note that machine learning approaches can generalize beyond their input parameters, making them less sensitive to changes in priors during training [17]. We assumed a mutation rate of 2.1x10^−10^ per site per generation, a generation time of one year [21] and a recombination rate of 8.25x10^-11^/site/generation estimated in the NEF cluster from a previous study [2]. We used boundaries for present effective population sizes between twice lower and five times higher their point estimates. We used a truncated exponential distribution for recombination rates with an average of 8.25x10^-11^ and a maximum value of 1.8x10^-9^. We conditioned on sweep completion occurring between 500,000 generations ago and shortly before northern clusters diverged from each other, around 150,000 generations ago. We allowed for variable gene flow between northern populations after their split (0<4*N_0_m*<0.5 with *N_0_*=1,475,000, the current effective population size of the GA cluster). Categorization of 30kb windows as sweeps, linked or neutral was then performed on the actual dataset, removing scaffolds shorter than 330 kb and sex-linked regions. diploS/HIC assigns a probability to each window for being a sweep, linked to a sweep, or neutral. We took advantage of this probability of assignation to improve the false positive rate by retaining candidate windows classified as sweeps only if they displayed a probability < 10% of being neutral.

**Genome scan using LSD**

To detect genomic regions displaying signatures of positive selection in the branch leading to northern clusters, we used the LSD algorithm [22]. This method compares the levels of exclusively shared differences between populations across genomic windows. The LSD score should be maximized when the local tree displays evidence for recent coalescence within the focal populations and high differentiation compared to the rest of the genome. Individuals were first phased using BEAGLE v4.0 with default options [23]. Haplotypes trees were computed over non-overlapping 1,250 SNPs windows with PhyML v3.1 [24] using the script phyml_sliding_windows.py available at <https://github.com/simonhmartin/genomics_general>. We chose to use 1,250 SNPs windows instead of windows of fixed lengths to get an equivalent amount of information across windows and increase resolution in regions of high recombination where SNP density and diversity is higher. The LSD algorithm was run over the 5 genetic clusters identified in the green anole (Figure 1, [2], Sup. Figure 1 in this study), using the branch leading to the two northern clades CA and GA as the focal branch in which to detect selection (CA and GA correspond to branches A1 and A2 in the description of the method available at <https://bitbucket.org/plibrado/LSD>) while the Floridian clade NEF was used as the sister branch to the ancestor of northern clades. The trees were rerooted by the algorithm using *Anolis allisoni* as an outgroup. We considered the windows ranking in the top 1000 LSD scores as candidates for positive selection and extracted overlapping genes (Ensembl release 90, genome version: AnoCar2.0) using the intersect function in bedtools v2.25.0 [25].

**Genome scan using BAYPASS**

We used the approach implemented in BAYPASS [26] to detect SNPs displaying high differentiation in northern populations. Overall divergence at each locus was first characterized using the *X^T^X* statistics, which is a measure of adaptive differentiation corrected for population structure and demography. Briefly, BAYPASS estimates a variance-covariance matrix reflecting correlations between allele frequencies across populations, a description that can incorporate admixture events and gene flow. This matrix is then used to correct differentiation statistics. BAYPASS offers the option to estimate an empirical Bayesian p-value (*eBPis*) which can be seen as the support for a non-random association between alleles and specific populations. We computed *eBPis* over the top 5% *X^T^X* outliers to determine their level of association with northern populations. BAYPASS was run using default parameters under the core model. We considered a SNP as a candidate for selection in northern populations when belonging to the top 1% *eBPis*. We divided the genome in 5kb windows and retained those with at least three outlier SNPs as candidates for positive selection. Genes overlapping these windows were extracted using bedtools v2.25.0.

**Overlap between scans of positive selection**

To evaluate the overlap and consistency between these methods, we examined the distribution of statistics for selection in regions classified as sweeps or neutral by diploS/HIC. To facilitate comparisons between LSD and *eBPis*, we computed the median *eBPis* score over the same windows as LSD. Candidate windows for selection (hard and soft sweeps combined) harbored an excess of high LSD and median *eBPis* scores compared to windows classified as neutral, the effect being particularly clear for the LSD statistics (see Figure 3 in [1]). Based on these distributions, we extracted a set of windows with consistent signals of selection. These 1,250 SNPs windows had a minimum overlap of 50% with windows classified as selected by diploS/HIC and belonged to the top 10% for median *eBPis* and LSD scores.
