## Supplementary figures and images for "Disentangling the determinants of transposable elements dynamics in vertebrate genomes using empirical evidences and simulations"

### Sup. Figure 1

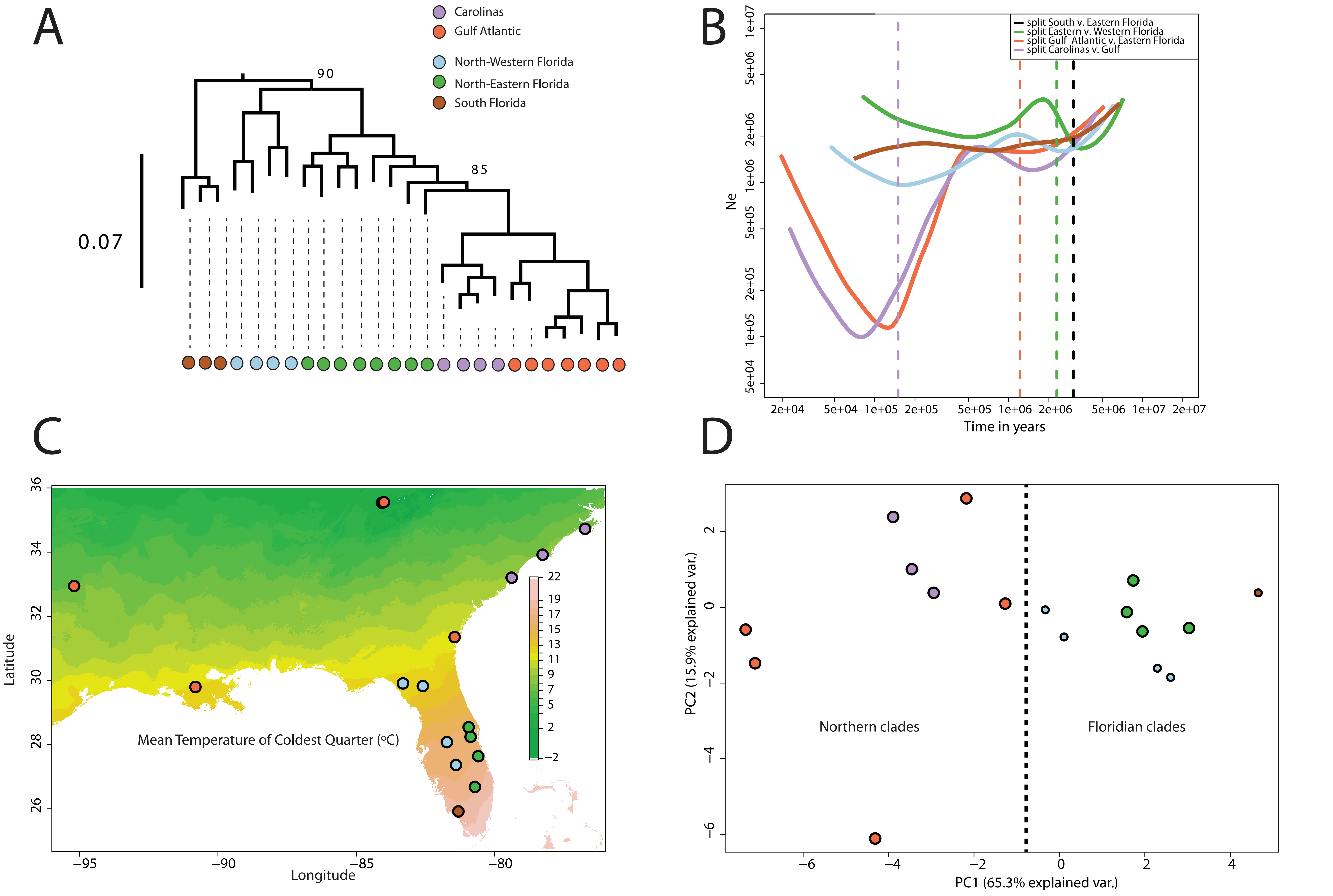

### Sup. Figure 2-5

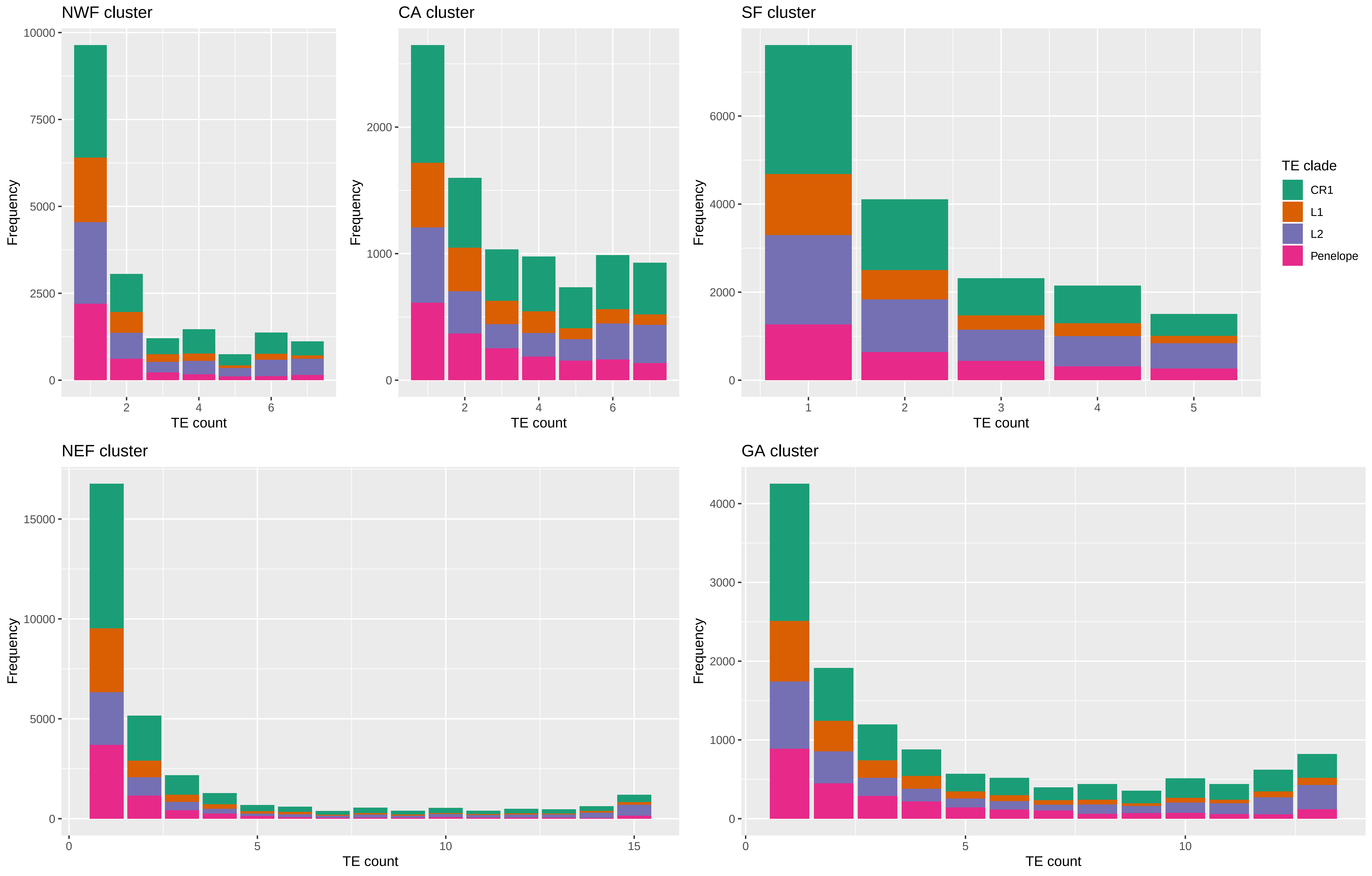

### Sup. Figure 2-5

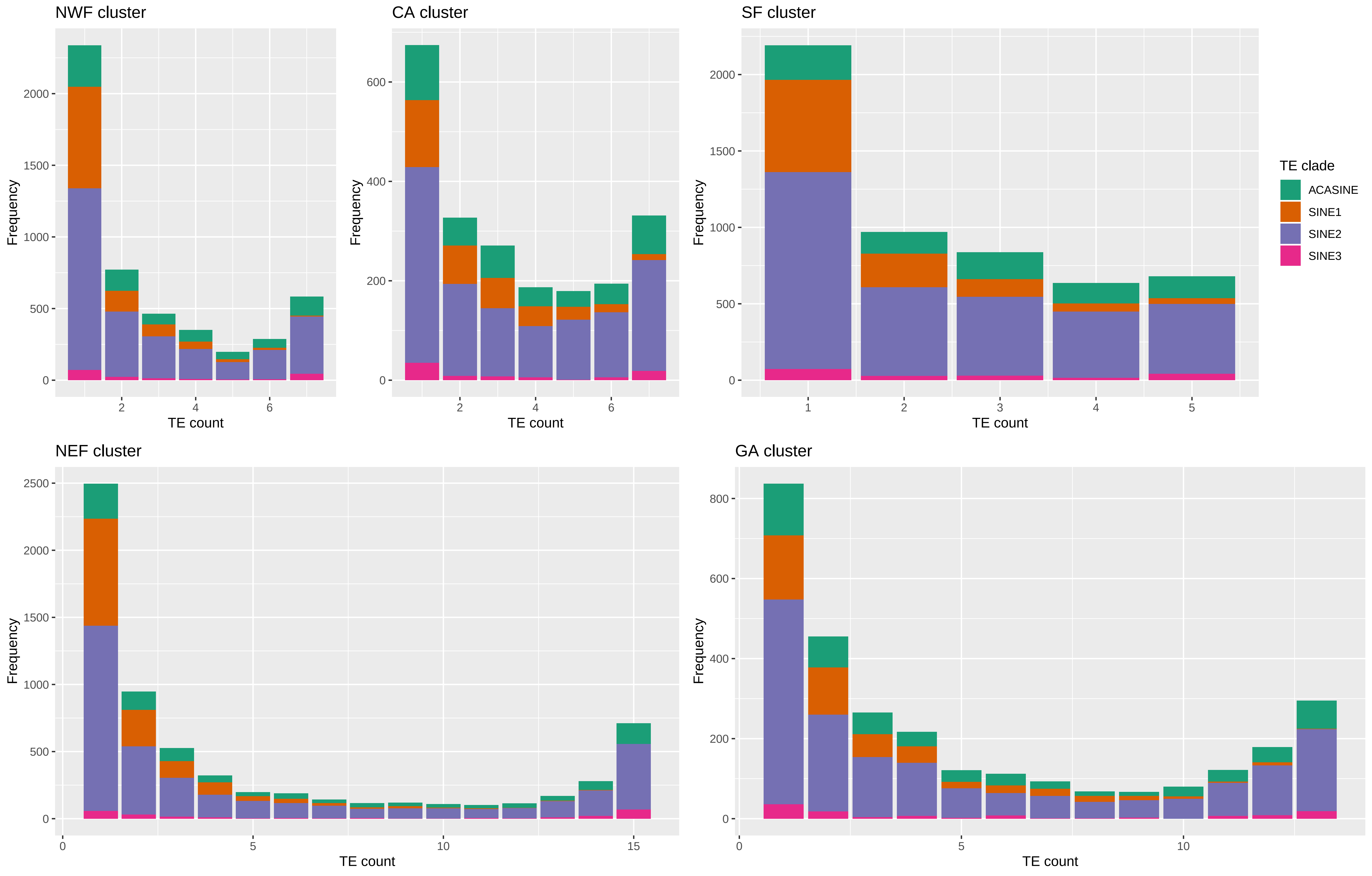

### Sup. Figure 2-5

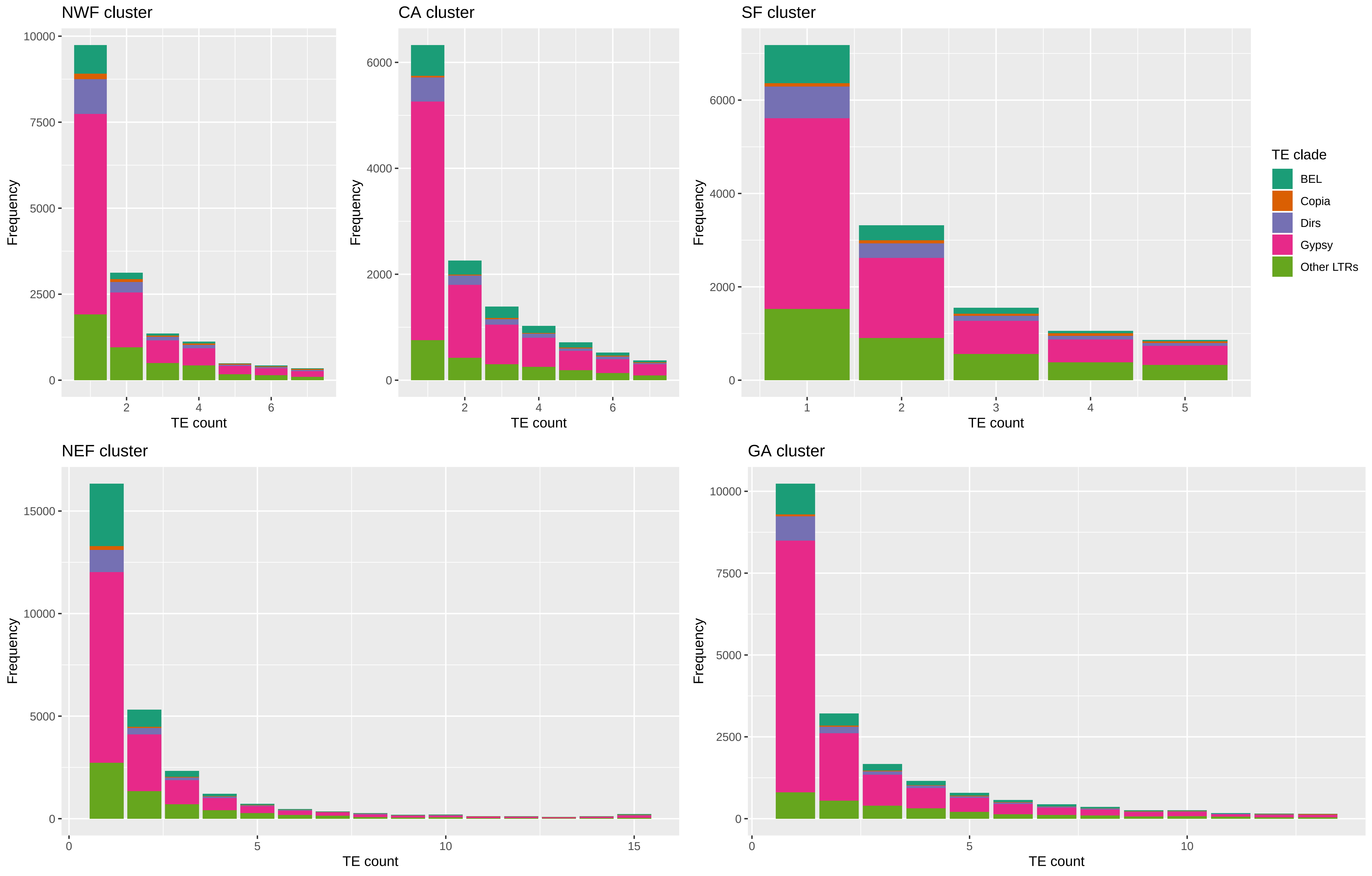

### Sup. Figure 2-5

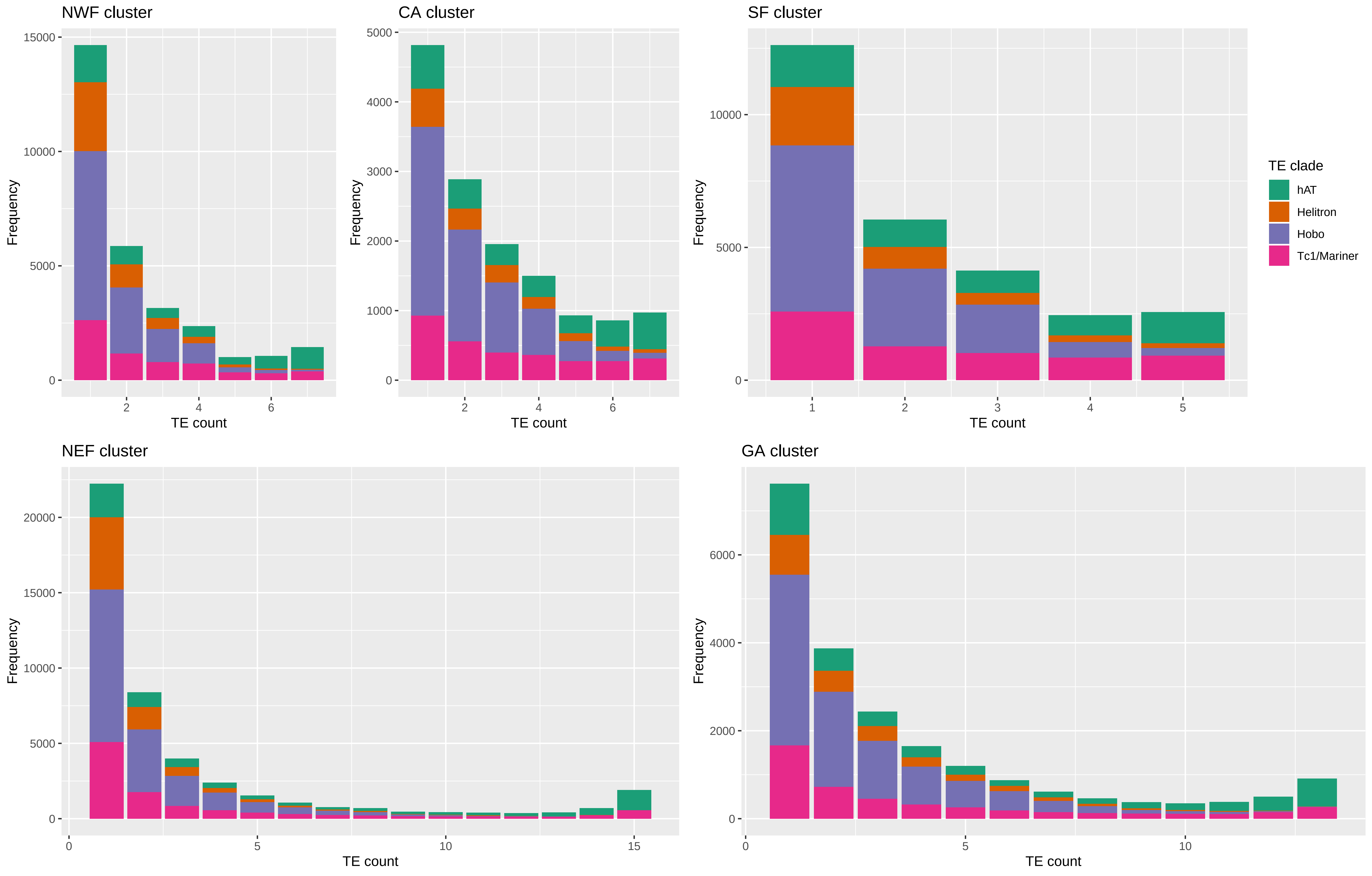

### Sup. Figure 6

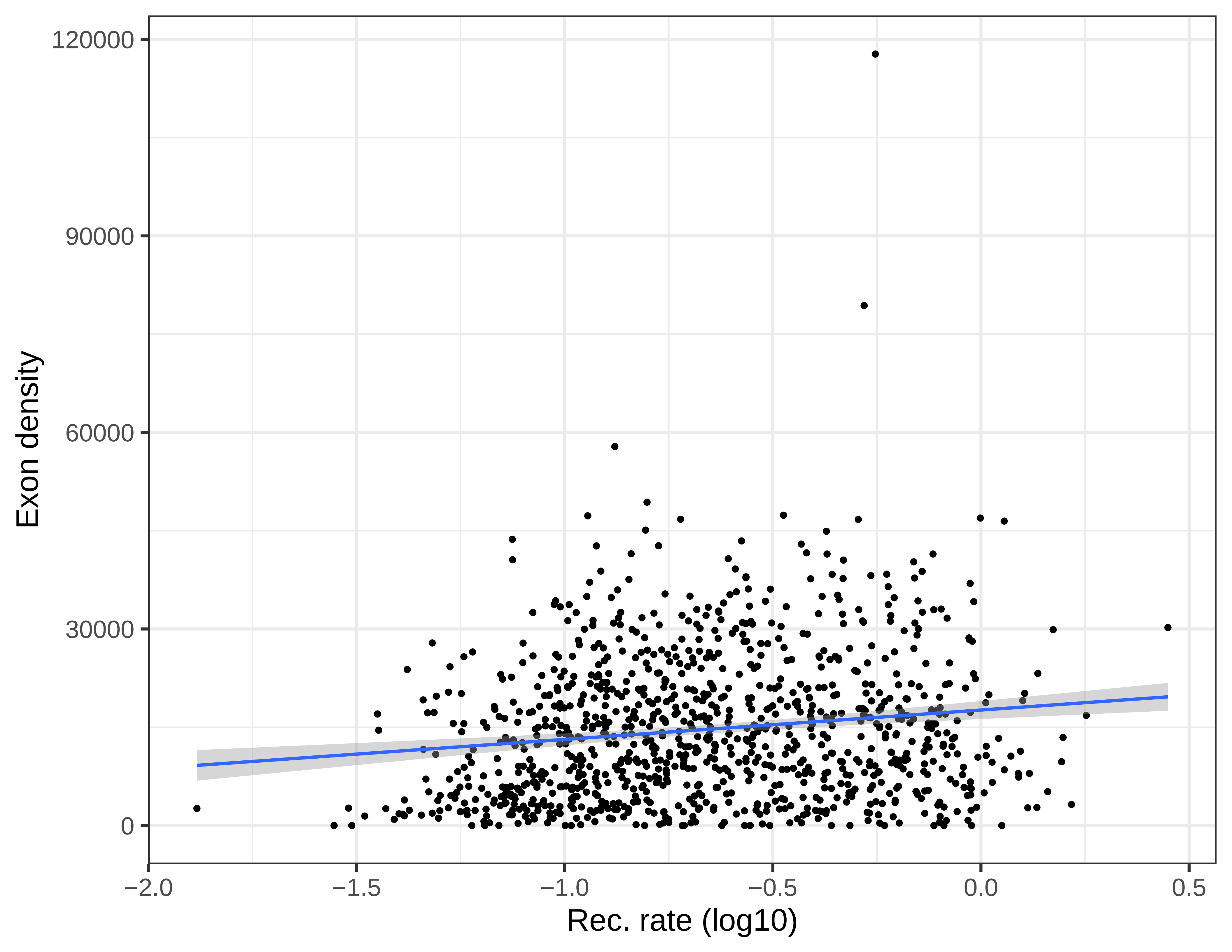
