## Supplementary material for "Disentangling the determinants of transposable elements dynamics in vertebrate genomes using empirical evidences and simulations": Sup. Figure 7

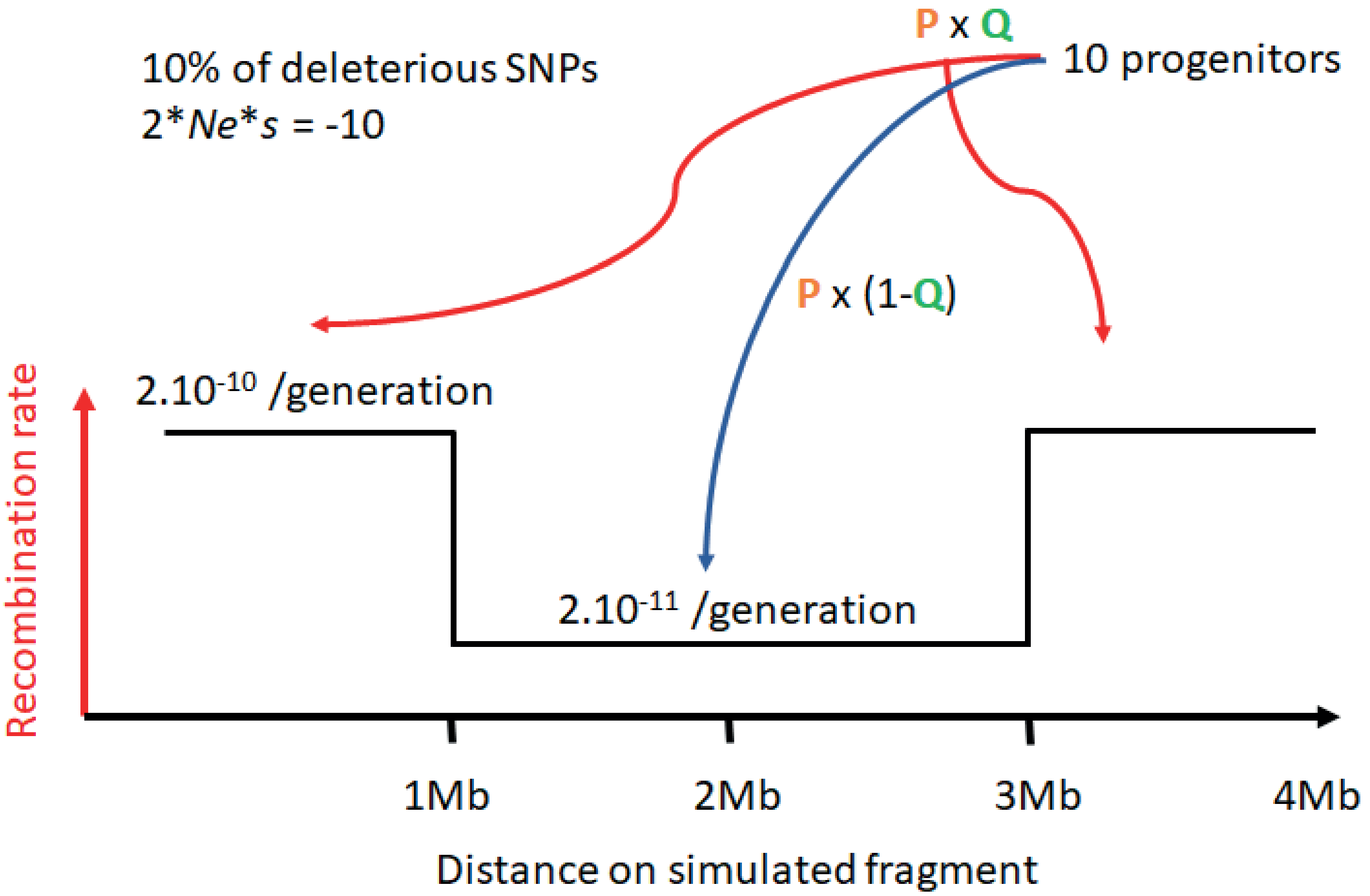


**Jumping probability (genome-wide):**

**$P=1.10^{-3}$ (constant transposition) or $1.10^{-1}$ (burst for 100,000 years)**

**Preferential insertion factor Q=0.5 or 0.7**
